# Estimates of habitat selection reveal distinct habitat associations across life-stages in three coral-reef damselfish

**DOI:** 10.64898/2026.03.26.714170

**Authors:** Gabriele Sciamma, Eric P. Fakan, Andrew S. Hoey

## Abstract

Understanding habitat association of animals and how they change through ontogeny is critical to predict the likely effects of habitat change on populations. We investigated how fine scale habitat associations of three common coral reef damselfish species changed among life-stages on reefs surrounding Lizard Island, northern Great Barrier Reef. All three species showed distinct habitat selection at settlement, however the degree to which these initial associations changed through ontogeny were species specific. *Pomacentrus amboinensis* associated with sandy areas throughout all life-stages; *Pomacentrus chrysurus* settled to areas with high cover of sand and rubble, but displayed no clear habitat preferences as juveniles or adults. *Pomacentrus moluccensis* settled to areas with high cover of fine branching corals before shifting to areas with relatively high cover of soft corals as adults. We also compared two different approaches to estimate habitat selection; one that quantified the benthic composition within the approximate home range of individuals versus a more widely used approach of recording a single point underneath the focal individual when they were first observed. Although results were broadly similar, the benthic composition approach revealed details that was overlooked using the single point method. Decreases in the availability of any of these preferred benthic habitats may adversely affect future populations, therefore understanding habitat associations and their transitions among life stages will be crucial in predicting future reef fish communities under ongoing coral loss and habitat change. This will require to systematically study a broader range of species, integrating relevant spatial and temporal scales.

## 1. INTRODUCTION

Habitat selection is a key component of an organisms’ ecology as it affects the way they interact with their environment and co-occurring species (Manly et al. 2004). Habitat selection, the non-random occupation of habitat, modulates biotic and abiotic interactions, influences the fitness of individuals and the distribution and dynamics of populations (Boyce & McDonald 1999, Morris 2003). Quantifying a species habitat preferences, and how these preferences differ among life stages is important to understand how habitat availability and future disturbances may shape populations and assemblages (e.g. Safi & Kerth 2004, Pratchett et al. 2012).

Tropical coral reefs are heterogeneous environments hosting approximately one third of all marine fish species (Reaka 1997, Bouchet & Duarte 2006). Coral reefs feature many microhabitats, allowing species to coexist through niche partitioning and differential habitat occupation (Wilson et al. 2007, Messmer et al. 2011). Because habitat-forming hard corals are fundamental in maintaining biodiversity (Wilson et al. 2007, Graham & Nash 2013, Coker et al. 2014), recent declines in the cover of live corals on reefs globally (Gardner et al. 2003, De’Ath et al. 2012, Hughes et al. 2017, Eakin et al. 2026) have led to concerns regarding the persistence of reef fish populations (Pratchett et al. 2008, Hoey et al. 2016). Notably, fish species that are dependent on live corals for food and/or habitat are the first and generally most adversely affected by reductions in coral cover (Munday 2004, Wilson et al. 2006, Pratchett et al. 2012). Yet, fish species that appear to have no direct reliance on live corals, at least as adults, have also been negatively impacted by the declines in coral cover (e.g. Booth & Beretta 2002, Pratchett et al. 2015). The mechanisms for such declines are not known and may be related to the effects of coral loss on their preferred prey items, or the use of live corals at different life stages. Indeed, about 39% of the fish species associating with live corals only do so at juvenile stage (Coker et al. 2014). Likewise, up to 65% of local reef fishes of Kimbe Bay (PNG) preferentially settle on live corals despite only 11% of species exhibiting obligate association with them as adults (Jones et al. 2004). These differences in the reliance on live coral among life stages highlight the ubiquity of ontogenetic habitat shifts in reef fishes and the importance of understanding these shifts to predict the effects of habitat change on fish populations. Large-scale coral mortality and the degradation of habitats may indeed affect reef fish species through stage-specific effects, such as settlement failure (Jones et al. 2004, Feary et al. 2007, Pratchett et al. 2008), and/or post-settlement processes that limit the number of individuals entering juvenile or adult life stages (Finn & Kingsford 1996, McCormick & Hoey 2004, Almany & Webster 2006).

Ontogenetic habitat shifts are widespread across the animal kingdom, where individuals balance the risks of predation with the acquisition of food as they grow (Werner & Gilliam 1984, Snover 2008). While great emphasis has been given to selection of habitats by coral reef fishes at settlement (Montgomery et al. 2001, McCormick et al. 2002), relatively few studies have quantified how these habitat associations change through ontogeny, despite ontogenetic habitat shifts being relatively common among reef fish (Eagle et al. 2001, Mellin et al. 2007, Clark & Russ 2012, Giffin et al. 2019, Valenzuela et al. 2021). In damselfish (f. Pomacentridae), one of the most extensively studied groups in terms of habitat requirements and associations (Pratchett et al. 2016), quantifications of ontogenetic shifts in habitat associations after settlement are lacking (for exceptions see Lirman 1994, McCormick & Makey 1997, Komyakova et al. 2019). Moreover, traditional quantifications of habitat associations by damselfish have used point observations of individuals, whereby habitat use is recorded as the benthic habitat directly underneath an observed individual. However, individuals use many of the available microhabitats within their home ranges (Streit et al. 2021). A comprehensive understanding of patterns of habitat associations requires an integration of the habitat composition within an individual’s home range compared to the available habitats, and how this varies among life stages. Achieving such understanding will increase our ability to predict differential impacts of habitat change on fish populations and reconcile observations which depart from theoretical predictions (Wismer et al. 2019a, 2019b).

We investigated habitat associations of three common coral reef damselfish species and how these associations vary with ontogeny. In particular, we tested how the selection of habitats changes among life stages (recently-settled, juvenile, and adult) and among species. Additionally, we compared estimates of habitat selectivity for recently-settled individuals of the three species using the single-point method (i.e., benthos directly under an observed individual) versus the benthic composition within an approximated home range (area method).

### 2. METHODOLOGY

#### 2.1 Study species and site

Three common damselfish species that differ in their habitat associations as adults were selected for this study: (i) *Pomacentrus amboinensis*, a ‘facultative’ coral-dweller (Pratchett et al. 2016); (ii) *Pomacentrus chrysurus* a habitat generalist that commonly inhabits rubble (Fakan et al. 2024) and (iii) *Pomacentrus moluccensis* which is considered an ‘obligate’ coral-dweller (Pratchett et al. 2012). Data were collected at Lizard Island, a high island located on the mid-shelf in the northern Great Barrier Reef (GBR; 14°40′ S, 145°28′E) during October - November 2021. Two sampling sites (each approx. 15 x 25 m) were selected on the sheltered south-west aspect of the island. The two sites were on the edge of shallow reefs characterized by a sandy reef base at ∼4m depth, a gentle reef slope up to a reef crest at ∼1m depth, and an extended reef flat.

#### 2.2 Sampling procedure

The two sites were systematically and thoroughly searched for newly-settled damselfish each day for two weeks (one week either side of the new moon) in both October and November 2021. This period coincides with the peak settlement period for reef fishes on the northern GBR (Meekan et al. 1993). The daily surveys allowed newly-settled damselfish of the three target species to be detected and sampled in the first 24-hours following settlement, thereby ensuring the detection of habitat associations at settlement that may be lost as a result of environmental filtering due to post-settlement mortality or movement. Juveniles and adults were classified based on body length (Table S1) and behaviour. A total of 395 individuals were sampled across the three species and three life stages (recently-settled, juveniles and adults) and two sites (Table S1). A minimum of 30 individuals per life stage were sampled in each species.

Recently-settled and juvenile fish were captured using a 10% clove oil:ethanol solution and hand-nets, transferred to a plastic bag where they were laterally photographed against grid paper (0.5 cm grid) *in situ*. Each captured individual was marked with a subcutaneous elastomer tag (Hoey and McCormick 2006) at the base of the dorsal fin to avoid resampling of individuals. Total length of recently-settled fish was recorded from the lateral photograph using ImageJ, and total length of juveniles was recorded *in situ* using callipers. Adult fishes were not captured and total length was visually estimated to the nearest centimetre.

#### 2.3 Habitat associations

To quantify the composition of benthic habitat surrounding each individual fish, quadrats were placed on the substratum centred over the location of first observation of the focal individual, and photographed from directly above at a height of approximately 1m. Quadrat sizes were selected based on estimated home ranges of each life stage (0.75 x 0.75 m for recently-settled and juvenile fish, and 1 x 1 m for adult fish; Table S1). Each photoquadrat was analysed in photoQuad where the substratum directly under each individual was recorded (single-point method) and under a series of stratified random points (area-method; 100 points for recently-settled and juvenile fish, and 125 points for adult fish). An additional 35 randomly-placed control quadrats (0.75 x 0.75 m) were sampled at each site (n = 70 in total) and analysed through the same procedure to assess the availability of benthic habitats. The habitat type under each of the random points was identified and grouped into one of eight categories according to their physical and biological attributes: (i) pavement, (ii) rubble, (iii) sand, (iv) macroalgae, (v) soft corals, (vi) fine branching corals – thin branches forming tight spaces such as tabular, corymbose and caespitose colonies, accessible by small fishes (e.g., *Seriatopora hystrix*), (vii) coarse branching corals – thick branches forming large spaces such as arborescent and digitate colonies (e.g., *Acropora muricata, Acropora pulchra*), and (viii) mounding corals – massive, encrusting and columnar morphologies (i.e. *Porites*).

#### 2.4 Habitat selection

Benthic habitat composition for each individual fish was expressed as relative proportion of each habitat type within its photoquadrat. Selection for each of the eight benthic categories was quantified using the forage ratio (*w_i_*) calculated using the ‘dietr’ package in R environment as (Manly et al. 2004):

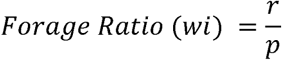

where *r* represents the proportion of the habitat utilized by an individual and *p* the proportion of the same habitat available at the study site. Values of *w_i_* below 1 indicated avoidance; for *w_i_*equalling 1, the habitat type was neither avoided or selected, being used in proportion to its availability. *w_i_* values exceeding 1 indicated selectivity for the habitat.

The forage ratio was first used to determine habitat preferences for recruits: an average of forage ratios was calculated for each species and life-stage with associated Bonferroni adjusted 95% confidence intervals following Manly et al. (2004) formula:

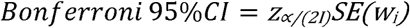

where *z*_∝/*(2I)*_ is the critical value of the standard normal distribution to an upper tail area of ∝*/2I* = 0.05 and *I* = the number of habitat types. Significance of habitat selection and avoidance was established when the 95% CI did not include *w_i_* = 1 (respectively higher and lower) (See supplementary material). Because of their negligible availability and scarce utilization by the study species, coarse branching corals were removed from the analysis, as their widespread absence across samples yielded a large number of zeros when calculating the forage ratios.

#### 2.5 Ontogenetic Habitat Shifts

To detect any difference in habitat associations among life stages, a one factor PERMANOVA based on Bray-Curtis distances was performed for each species, with life stage as a factor (4 levels: recruit, juvenile, adult, and control). For species returning significant models, pairwise comparisons were implemented among the three life stages and significance established using Bonferroni adjusted p-values. Significant differences in habitat between life-stages indicated ontogenetic habitat shifts. To visualize differences in habitat association among life-stages a Canonical Discriminant Analysis (CDA) was used. Vector plots were imposed to identify the critical habitat types driving significant differences in habitat associations among factors. All analysis were performed with R in RStudio environment.

### 3. RESULTS

#### 3.1 Habitat availability and habitat selection in recruits

The benthic composition of the study sites was characterized by a high cover of pavement (∼ 35%), moderate cover (14-16%) of hard corals (fine branching and mounding corals combined), soft coral as well as rubble, and low cover (4-11%) of algae and sand (Fig. 1a). Recently-settled *Pomacentrus amboinensis* showed strong positive selection for areas with high cover of sand (35% cover; *w_i_*= 3.40 ± 0.75 CI), avoided areas characterized by algae, pavement and rubble, and used hard and soft corals proportional to their availability (Fig. 1b, e). Recently-settled *Pomacentrus chrysurus* also selected for areas with high cover of sand (20% cover; *w_i_* = 1.90 ± 0.39 CI) and rubble (33% cover; *w_i_* = 1.98 ± 0.29 CI), and avoided soft coral, hard coral, and algae (Fig. 1c, f). In contrast, recently-settled *Pomacentrus moluccensis* selected areas with high cover of fine branching corals (16% cover; *w_i_* = 2.26 ± 0.38 CI) and avoided areas of algae (2% cover; *w_i_* = 0.44 ± 0.22 CI) and rubble (11% cover; *w_i_* = 0.63 ± 0.19 CI) (Fig. 1d, g). All other substrata were used in proportion to their availability (Fig. 1g).

**Figure 1.**
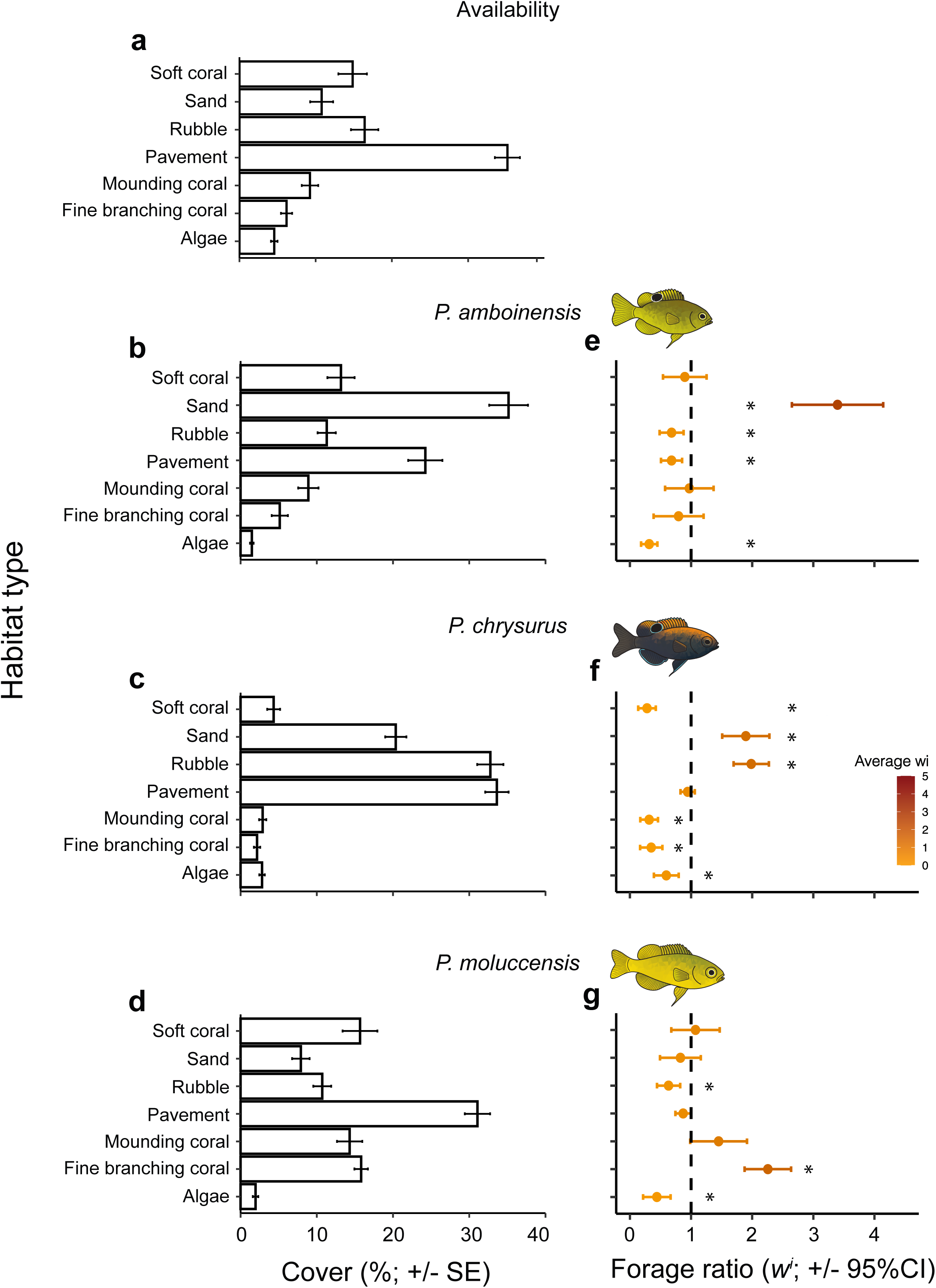
Availability (a), use (b-d) and selectivity (e-g) of benthic habitats by three species of recently-settled damselfish: (b,e) *P. amboinensis*, (c,f) *P. chrysurus*, (d,g) *P. moluccensis*. The availability and use of each substratum type is expressed as average percent cover ± 1 SE. Selectivity of substratum types by recently-settled fish is calculated as the forage ratio (*w*_i_), with the error bars showing Bonferroni corrected 95% confidence intervals. A forage ratio (– 95% CI) greater than one indicates positive selection, and a forage ratio (+95% CI) less than one indicates negative selection (or avoidance). If the 95% CI includes one, the substratum was used in proportion to its availability.

#### 3.2 Selectivity through ontogeny

The habitat associations of *P. amboinensis* were relatively consistent among life stages, with juvenile and adult *P. amboinensis* maintaining a positive selection for sand and avoidance of algae and pavement (Fig 2a, d). While recently-settled and juvenile *P. amboinensis* used fine branching coral habitats in proportion to its availability (Fig. 1e, 2a) adult conspecifics used fine branching corals less than expected based on its availability (*w_i_* = 0.43 ± 0.31 CI; Fig. 2d). The habitat associations of both *P. chrysurus* and *P. moluccensis* changed considerably among life stages. While recently-settled *P. chrysurus* selected for rubble and sand, juvenile conspecifics selected rubble (*w_i_* = 1.46 ± 0.43 CI) only, and adult *P. chrysurus* did not positively select any habitat type (Fig. 2b, e). All three life-stages of *P. chrysurus* consistently avoided hard coral habitat types (mounding and fine branching corals) whereas soft corals were avoided by recently-settled and adult individuals (*w_i_* = 0.67 ± 0.26; Fig. 2e), but not juveniles. Likewise, the positive selection for fine branching coral by recently-settled *P. moluccensis* was reduced in juvenile conspecifics (*w_i_* = 1.68 ± 0.62 CI), and adults showed no selection for this habitat type (Fig. 2c, f). Recently-settled and adult *P. moluccensis* used algae habitat proportionally less than its availability, while juveniles showed no selection or avoidance for algae (Fig. 2c, f).

**Figure 2.**
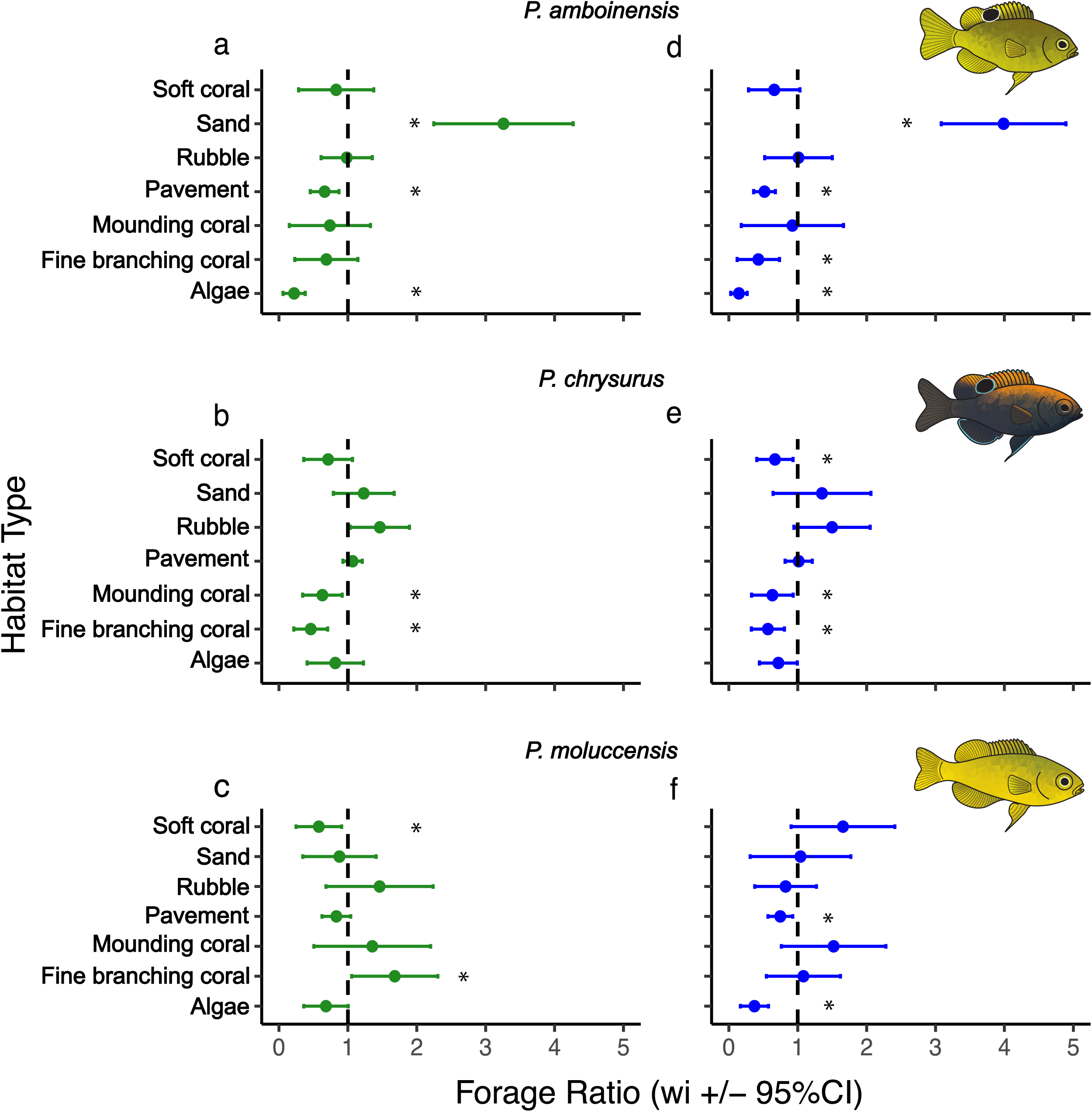
Selectivity of benthic habitats by juvenile (green) and adult (blue) life stages of the three study species: (a,d) *P. amboinensis*, (b,e) *P. chrysurus*, (c,f) *P. moluccensis*. Selectivity of substratum types is calculated using the forage ratio (*w_i_*) ± Bonferroni corrected 95% confidence intervals. A forage ratio (–95% CI) greater than one indicates positive selection, and a forage ratio (+95% CI) less than one indicates negative selection (or avoidance). If the 95% CI includes one, the substratum was used in proportion to its availability.

#### 3.3 Ontogenetic habitat shifts

Comparisons of the habitat composition of the three life stages of *P. amboinensis* and control plots showed that while all life stages associated with areas that had a higher abundance of sand compared to the control plots (PERMANOVA: p < 0.001, R^2^ = 0.20), there were no significant differences in habitat composition among life stages (Fig. 3a, Table S2).

**Figure 3.**
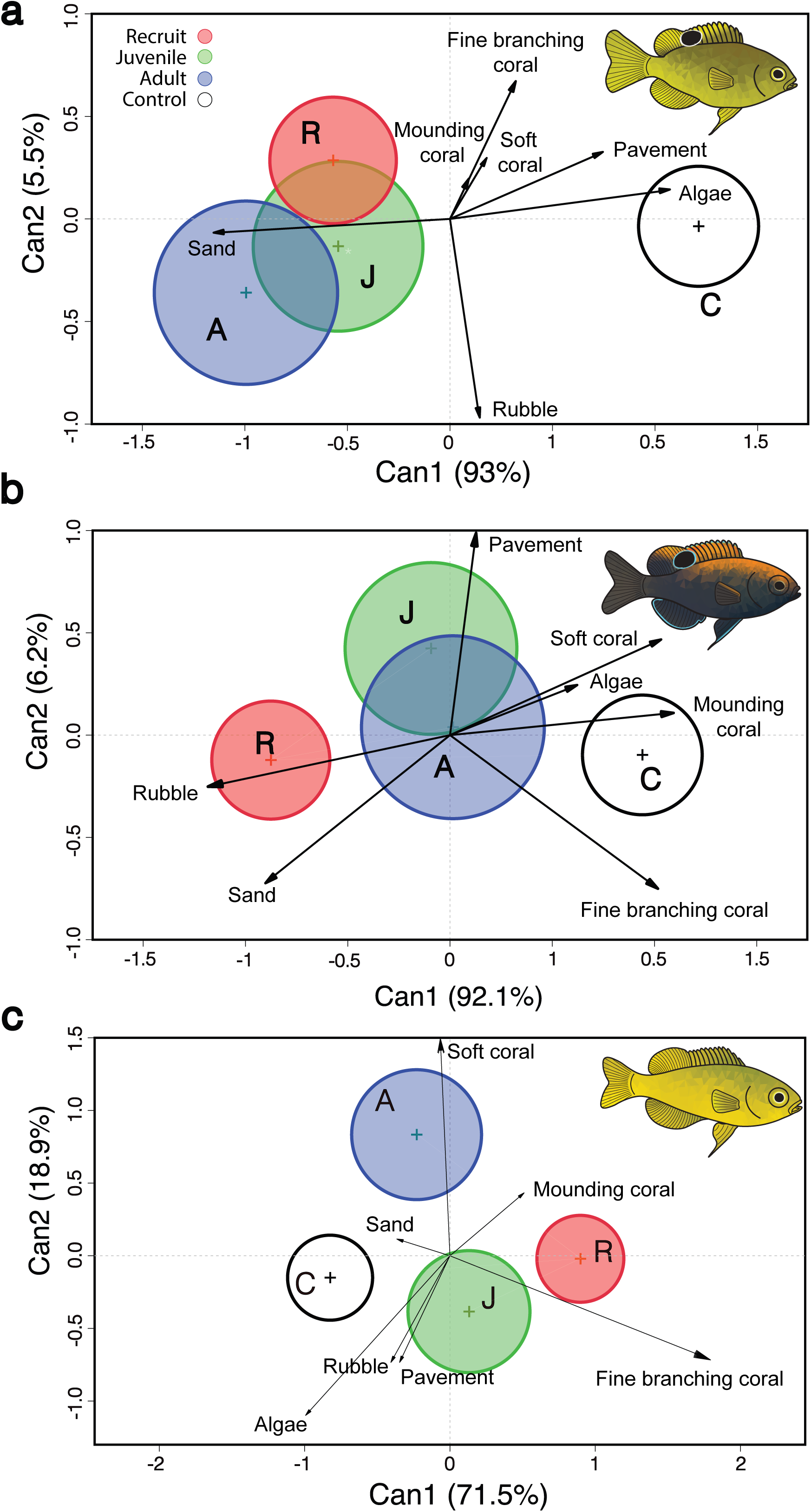
Canonical Discriminant Analyses (CDAs) of benthic habitat composition in (a) *P. amboinensis*, (b) *P. chrysurus* and (c) *P. moluccensis*. Each life-stage is represented by the 95% confidence ellipse of the canonical scores around the mean centroid. Each habitat type is represented by a vector, the length of which indicates its relative importance in separating samples according to life-stages.

There were, however, significant differences in the habitat associations of *P. chrysurus* among life stages (PERMANOVA: p < 0.001, R^2^ = 0.12). The habitats of recently-settled *P. chrysurus* differed from those of conspecific juvenile and adult life stages, and it was the only life stage to differ from the control plots (Table S2). The CDA showed that recently-settled *P. chrysurus* associated with areas with relatively high cover of sand and rubble, and low cover of live corals, whereas control plots were characterised by soft, mounding and fine branching corals as well as algae (Fig. 3b).

For *P. moluccensis* each of the three life stages exhibited distinct habitat associations (PERMANOVA: p < 0.001, R^2^ = 0.07; Table S2). The habitat associations of recently-settled *P. moluccensis* were characterised by a relatively high cover of fine branching coral, whereas conspecific juveniles were characterised by a relatively high cover of rubble, algae and pavement, and adult conspecifics were characterised by relatively high cover of soft corals (Fig. 3c). The habitat composition of the juvenile life-stage was the only one which did not significantly differ from habitat found in control plots (Table S2).

#### 3.4 Method comparison

Estimates of habitat selection (forage ratio) based on the multi-point (area method) and single-point data in recently-settled damselfish yielded broadly similar results (Fig. 4), with a few interesting differences. Both methods indicated that recently-settled *P. amboinensis* used coral habitat types in proportion to their availability and avoided algae and, to a lesser degree, rubble. However, the area method indicated selection for sand, whereas according to the single-point method this habitat was used in proportion to its availability (Fig. 4a). It also detected an avoidance of pavement, which was not evident using the single-point observations. In recently-settled *P. chrysurus*, the area method detected selection for rubble and sand, random utilization of pavement and avoidance of any other substratum type. Estimates based on the single-point observations confirmed selection for rubble and avoidance for soft corals but showed random utilization of all the other habitat types, including sand and hard corals (Fig. 4b). Recently-settled *P. moluccensis* selected fine branching corals while avoiding algae and rubble according to estimates of benthic area composition. Estimates from single point observation instead highlighted avoidance sand, rubble and pavement, use of algae according to availability and strong positive selection for fine branching corals (Fig. 4c). Interestingly, the estimated forage ratio using the area method (9.03 ± 1.84 CI) was nearly four-fold greater than that of the single point method (2.26 ± 0.60 CI; Fig. 4c).

**Figure 4.**
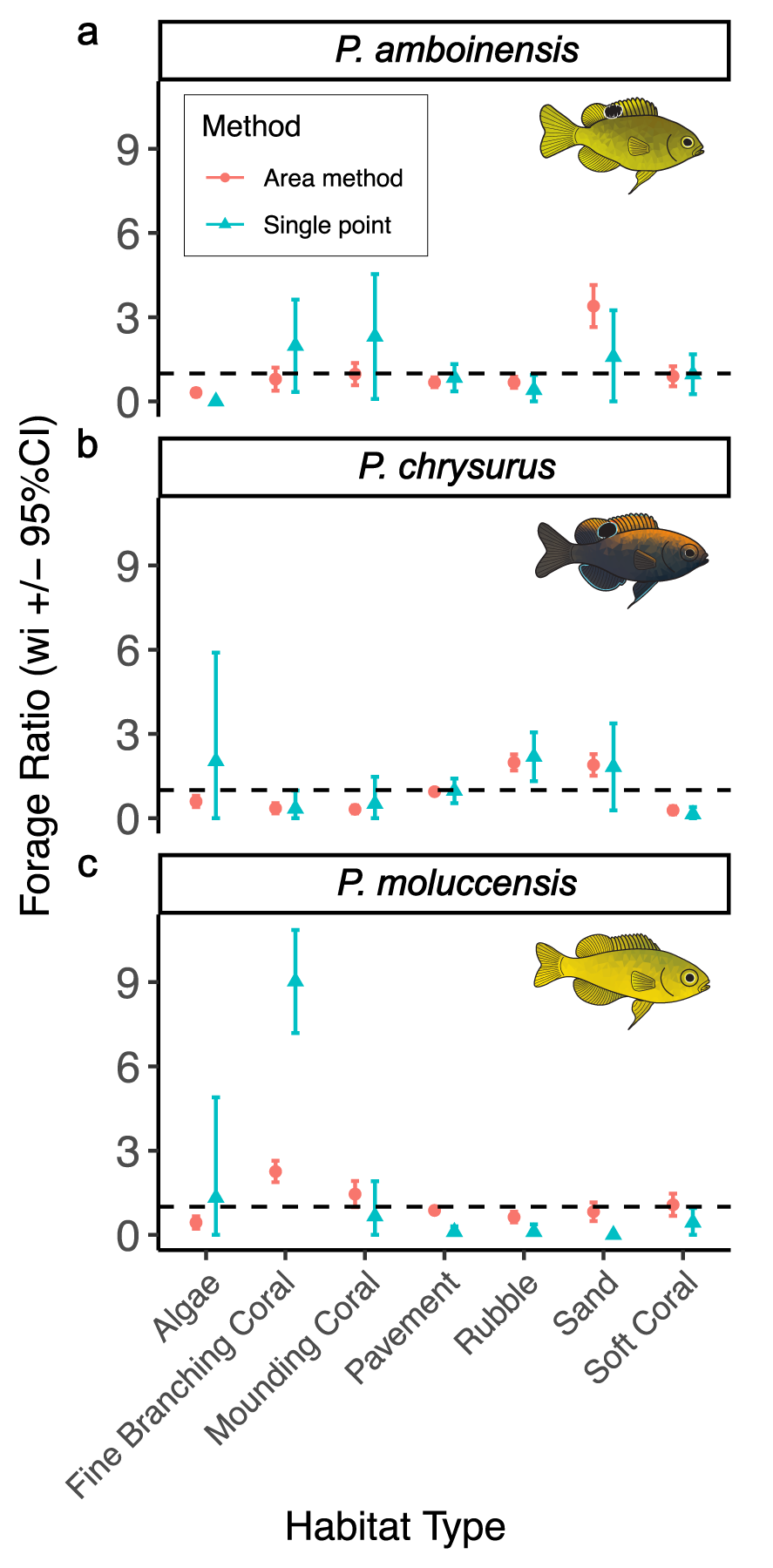
Comparison of habitat selection estimates (average forage ratio *w_i_*) using single-point (blue triangle) and area method (red circle) methods for each habitat type in the three study species at recruit stage. (a) *P. amboinensis,* (b) *P. chrysurus,* and (c) *P. moluccensis*.

### 4. DISCUSSION

The habitat associations of animals often reflect trade-offs between food availability, predation risk, and/or access to mates (Creel et al. 2005, Thomson et al. 2006, Grol et al. 2011). As such, habitat associations may change with ontogeny as the relative importance of predation risk, access to mates and/or energy requirements varies with body size and/or life stage (Werner & Gilliam 1984, Snover 2008). Coral reef damselfishes are highly site-attached, and as such are often assumed to settle directly into their juvenile and adult habitat (McCormick & Makey 1997, Lecchini & Galzin 2005, Komyakova et al. 2019). Here we showed that habitat associations of three species of damselfish are species and life-stage dependent. The three species had distinct habitat preferences shortly after settlement with recently-settled *Pomacentrus amboinensis* positively selecting areas with sand and avoiding algae, pavement, and rubble; *Pomacentrus moluccensis* selecting fine branching corals and also avoiding algae and pavement, and *Pomacentrus chrysurus* selecting rubble and sand and avoiding hard corals. The degree to which these initial habitat associations were maintained in later life stages also varied among the three species. While there was limited change in the habitat associations of *P. amboinensis* among life stages, *P. chrysurus* and *P. moluccensis* changed habitat associations among life stages, with recently-settled individuals of both species displaying greater selectivity (both positive and negative) of habitat types, than juvenile and adult conspecifics. These ontogenetic shifts in habitat associations could be related to selective post-settlement mortality (e.g., McCormick and Hoey 2004) and/or ontogenetic movements among habitat types (e.g., McCormick and Makey 1997). Importantly, the magnitude of these ontogenetic changes in habitat associations does not appear to be related to the degree of association with live corals; the greatest changes in habitat associations being observed in the obligate coral dweller *P. moluccensis* and the habitat generalist *P. chrysurus*, while the facultative coral dweller *P. amboinensis* displayed limited changes in habitat associations.

Settlement and the first few days post-settlement are a critical period in the life history of coral reef fish, with up to 75% of individuals being lost in the first five days post-settlement (Hoey & McCormick 2004, Almany & Webster 2006). Moreover, this high mortality can be selective, favouring individuals based on the characteristics of their habitat (McCormick and Hoey 2004, Blandford et al. 2023), as well as the local fish assemblage (Carr and Hixon 1995, Bonin et al. 2009) and their physiological state (Hoey and McCormick 2004, Dingeldein & White 2016, Fakan et al. 2024), thereby leading to differences in habitat associations among life stages. Although all three damselfish species are a similar size at settlement, and presumably similarly susceptible to predation, only one species (*P. moluccensis*) associated with a structurally complex habitat (i.e., fine branching coral), that would provide potential refugia from predation, at settlement (Coker et al. 2009). The other two species, *P. amboinensis* and *P. chrysurus,* settled to habitat types of low complexity, characterized by sand, and sand and rubble respectively. While the settlement of *P. moluccensis* to live branching corals and *P. chrysurus* to sand and rubble is consistent with previous field and aquaria studies (Ault and Johnson 1998, Ohman et al. 1998, Feary et al. 2007, Fakan et al. 2024), the association of recently-settled *P. amboinensis* with areas of high cover of sand is counter to previous aquaria studies that have shown this species to select structurally complex habitats, namely live and dead corals (Ohman et al. 1998, Feary et al. 2007). Field studies have, however, reported *P. amboinensis* to settle to a range of habitats including live and dead branching corals (Leis and Carson-Ewart 2002, McCormick and Hoey 2006, McCormick et al. 2010), rubble and sand (Ault and Johnson 1998, McCormick and Hoey 2004, Fakan et al. 2024), suggesting that any habitat preferences may be altered by other factors, such as the abundance of potential competitors (e.g., McCormick and Hoey 2004, Fakan et al. 2024) or preference for particular reef zones (e.g., reef edge: Leis and Carson-Ewart 2002).

Two of the three species of damselfish examined (*P. chrysurus* and *P. moluccensis*) underwent shifts in habitat association among life stages (cf. Komyakova et al. 2019), while there was limited change in the habitat associations of *P. amboinensis*. For both *P. chrysurus* and *P. moluccensis* the degree of habitat selectivity decreased with age, with recently-settled individuals of both species displaying greater selectivity (both positive and negative) of habitat types, than juvenile and adult conspecifics. These changes could be due to selective mortality, movement of individuals, and/or differences in the area sampled for each life stage. Differences in the area sampled between each life stage appears unlikely to have contributed to the observed ontogenetic shifts as the greatest difference was between recently-settled and juvenile fish, that were sampled using the same quadrat (i.e., 0.75 x 0.75m). While the relative importance of selective mortality and movement of individuals cannot be determined based on our results, previous research that tracked the fate of tagged individuals found that local fish assemblages had a greater effect on the survival of *P. moluccensis* and *P. chrysurus* in the 14 days following settlement (Fakan et al. 2024). For example, the presence and abundance of congeneric juveniles (< 2cm TL) had the greatest effect (∼30% reduction in mortality) on the survival of newly-settled *P. moluccensis.* The lack of an ontogenetic shift in the habitat association of *P. amboinensis* is consistent with a previous study (McCormick and Makey 1997), although somewhat counter to studies that have reported early post-settlement mortality to be related to the settlement habitat. Notably, McCormick & Hoey (2004) found mortality in the first 10 days post-settlement was greatest for *P. amboinensis* that had settled to sand and rubble, intermediate for those that had settled to *Pocillopora* and lowest for those that had settled to open branching corals. Similarly, Fakan et al. (2024) found that local rugosity had the greatest effect on the survival of *P. amboinensis* over the first 14 days post-settlement. The lack of ontogenetic shift in habitat association of *P. amboinensis* in the present study despite habitat-selective early post-settlement mortality (McCormick and Hoey 2004, Fakan et al. 2024) suggests that individuals may move among habitats and/or be subject to further selective mortality from 2-weeks post-settlement until they enter the juvenile population.

It should be noted that some of the differences in reported habitat associations and selectivity among studies may be related to the specific question being addressed and the methods used, in particular whether selectivity was based on the immediate habitat on which individuals were first observed or the broader habitat within their home range (i.e., single-point vs area methods). Indeed, comparison of the single-point and area method in the present study show that the positive selection for sand by recently-settled *P. amboinensis* using the area method was not evident using the single-point method. Moreover, the single-point method suggested that recently-settled *P. amboinensis* avoided algae, but used all other habitats in proportion to their availability. Collectively, these results indicate that *P. amboinensis* settles to a range of benthic habitats in areas of high sand cover, and is consistent with observations of *P. amboinensis* larvae settling near the reef edge (Booth 2002, Leis and Carson-Ewart 2002).

Similarly, adults of the humbug damselfish *Dascyllus aruanus* and juveniles of the bar-cheek coral trout *Plectropomus maculatus* have been reported to select branching corals surrounded by sand (Hoolbrook et al. 1999, Wen et al. 2013). This preference for areas of high sand cover has been suggested to be related to the availability of dietary resources (e.g., micro-crustacea or emergent plankton; Ault and Johnson 1998) and warrants further investigation. Similarly, comparisons of the single-point and area-based methods for assessing habitat selectivity revealed differences for *P. chrysurus* and *P. moluccensis*. Specifically, the single point method resulted in a 4-fold increase in the selectivity (i.e., forage ratio) of fine branching corals for recently-settled *P. moluccensis* compared to the area-based method, as well as the avoidance of sand and pavement that were both used in proportion to availability using the area-based method. These examples highlight the importance of integrating different spatial scales in studies of habitat association, especially if we are to understand likely changes to fish populations and communities under ongoing climate change and habitat degradation.

The three focal damselfish species in this study are often categorised based on their habitat preferences, with *P. amboinensis* typically referred to as ‘facultative coral-dweller’, *P. moluccensis* as on ‘obligate coral-dweller’ and *P. chrysurus* as a habitat generalist or rubble specialist (Low 1971, Pratchett et al. 2016). While these categories may broadly hold for *P. moluccensis* that showed a positive selection for fine branching corals at recently-settled and juvenile (but not adult) life stages, and *P. chrysurus* that selected rubble in each of the three life stages, there was no evidence that *P. amboinensis* selected for corals at any life stage. Rather *P. amboinensis* appeared to be a habitat generalist.

### 5. CONCLUSION

The present study showed that three congeneric damselfish species possess distinct habitat associations within the area they settle and live. Selection was found for habitat types within the individuals’ home-ranges which differed from previous findings based on single point observations. Furthermore, these non-random associations were limited to distinct life-stages for *Pomacentrus chrysurus* and *Pomacentrus moluccensis*. Thus, the integration of methods applying different spatial and temporal scales will likely provide fundamental insights into patterns of habitat utilization and the overall role played by benthic composition in accommodating fish populations. Because selection of certain benthic areas can vary at different life-stages, we conclude that changes in the benthic community composition may affect species through stage-specific effects. The ubiquity of habitat selection at settlement suggest that selective pressures may be greatest for this life stage and consequently may be particularly vulnerable to habitat change. Future studies on habitat requirements should integrate observations at different spatial scales and life stages to provide a deeper understanding of the interaction between fish species and changing coral reefs.

## Acknowledgements

We thank the staff of Lizard Island Research Station for the support to fieldwork operations.

## Author contributions

All authors conceptualized the study and designed the methods. EPF collected and extracted the data. GS analysed the data and wrote the first draft of the manuscript. The manuscript for submission was substantially revised by all the authors.

## Declarations

### Conflict of interest

The authors declare no competing interests.

### Ethical approval

All research was carried out under approval of the James Cook University animal ethics committee (permit: A2683) and according to the University’s animal ethics guidelines.

### Funding

Funding was provided by the Australian Research Council Centre of Excellence for Coral Reef Studies (ASH), the Lizard Island Reef Research Foundation (Gough Fellowship; EPF), and the Australian Museum’s Lizard Island Research Station (EPF).

### Data Availability

Dataset, codes and the computational environment in which the analyses were run are publicly available on GitHub (https://github.com/sciammag/sciamma_etal2026_habitatselection).

## Notes

https://github.com/sciammag/sciamma_etal2026_habitatselection

### Competing Interest Statement

The authors have declared no competing interest.

