## Supplementary material for "Estimates of habitat selection reveal distinct habitat associations across life-stages in three coral-reef damselfish": supp_material.docx

**Supplemental materials**

**Table S1.** Summary table showing the sample size (*n*), mean total length with its standard error (se), and the quadrat size for each life stage of the three species as well as the control samples.

| **Species** | **Life Stage** | **n** | **Mean TL (cm)** | **se (TL)** | **Quadrat Size (m)** |
| --- | --- | --- | --- | --- | --- |
| P. amboinensis | Recruit | 63 | 1.72 | 0.01 | 0.75x0.75 |
|  | Juvenile | 35 | 3.99 | 0.12 | 0.75x0.75 |
|  | Adult | 30 | 7.37 | 0.14 | 1x1 |
| P. chrysurus | Recruit | 72 | 1.89 | 0.01 | 0.75x0.75 |
|  | Juvenile | 34 | 4.05 | 0.09 | 0.75x0.75 |
|  | Adult | 30 | 7.00 | 0.10 | 1x1 |
| P. moluccensis | Recruit | 67 | 1.55 | 0.01 | 0.75x0.75 |
|  | Juvenile | 34 | 3.90 | 0.08 | 0.75x0.75 |
|  | Adult | 30 | 6.54 | 0.16 | 1x1 |
| Control | - | 70 | - | - | 0.75x0.75 |

**Table S2.** Summary table, comprehensive of R^2^ and adjusted p-values, of PERMANOVA pairwise comparisons between the three life-stages (recruit, juvenile, adult) and control plots for each species. Significant comparisons (p-adjusted < 0.05) are indicated by an asterisk (*).

| **Species** | **Pairs** | **Global R^2^** | **p-adjusted** |
| --- | --- | --- | --- |
| *P. amboinensis* | recruit vs control* | 0.152 | 0.0006 |
|  | juvenile vs control* | 0.137 | 0.0006 |
|  | adult vs control* | 0.235 | 0.0006 |
|  | recruit vs juvenile | 0.001 | 1 |
|  | juvenile vs adult | 0.022 | 0.9545 |
|  | recruit vs adult | 0.028 | 0.1626 |
| *P. chrysurus* | recruit vs control* | 0.17 | 0.0006 |
|  | juvenile vs control | 0.031 | 0.2010 |
|  | adult vs control | 0.027 | 0.9287 |
|  | recruit vs juvenile* | 0.071 | 0.0006 |
|  | juvenile vs adult | 0 | 1 |
|  | recruit vs adult* | 0.067 | 0.0036 |
| *P. moluccensis* | recruit vs control* | 0.066 | 0.0006 |
|  | juvenile vs control | 0.029 | 0.0870 |
|  | adult vs control* | 0.041 | 0.0144 |
|  | recruit vs juvenile* | 0.049 | 0.0210 |
|  | juvenile vs adult* | 0.068 | 0.0060 |
|  | recruit vs adult* | 0.051 | 0.0030 |
